## Supplementary material for "Phenotypic and genomic diversification in complex carbohydrate degrading human gut bacteria": Compressed data folder containing 4 pangenome maps and legend: Pudlo_Urs_Figure_S6_LEGEND.pdf

Pangenome genes:

- susC-like TonB dependent transporter
- susD-like
- susR
- susE
- susF
- Glycoside Hydrolase/Carbohydrate Binding Moiety
- glycosyl Transferase
- HTCS regulator
- ECF sigma regulator
- Anti sigma regulator
- ORF present in a PUL
- Transposase/Recombinase/Integrase/Xer-D family
- Tra Operon
- LacL Operon
- AsnC
- Gent/Tet Resistance
- Conserved PUL gene
- Variable PUL gene

Pangenome loci (PULs, conserved, non-conserved, etc.) :

- Variable genomic region
- Strain variable PULs
- Conserved PULs
- Conserved PULs (highly divergent ORFs)
- Strain-varibale PULS (divergent ORFs)
- Capsular poysaccharide synthesis
