## Supplementary figures and images for "Phenotypic and genomic diversification in complex carbohydrate degrading human gut bacteria"

### Pudlo_Urs_et_al_Fig_S6a_pangenome1.pdf

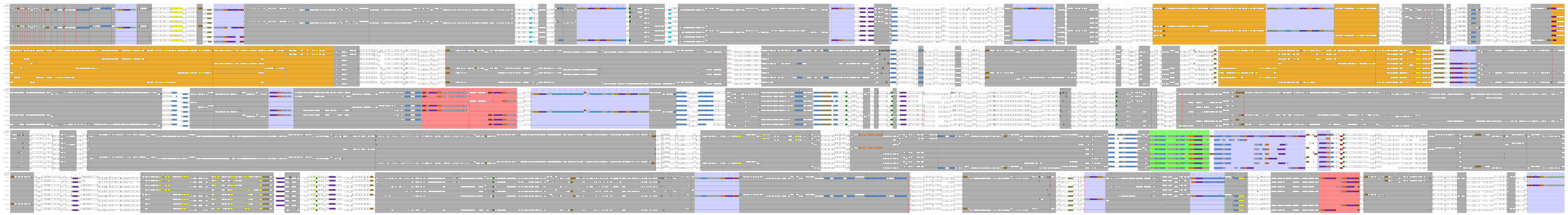

### Pudlo_Urs_et_al_Fig_S6b_pangenome2.pdf

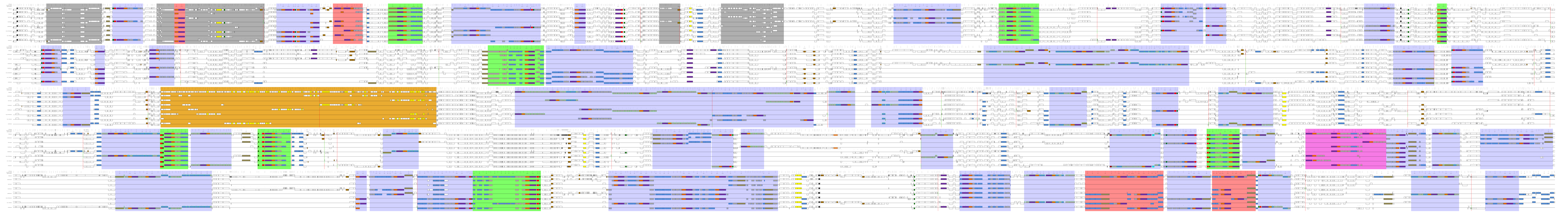

### Pudlo_Urs_et_al_Fig_s6c_pangenome3.pdf

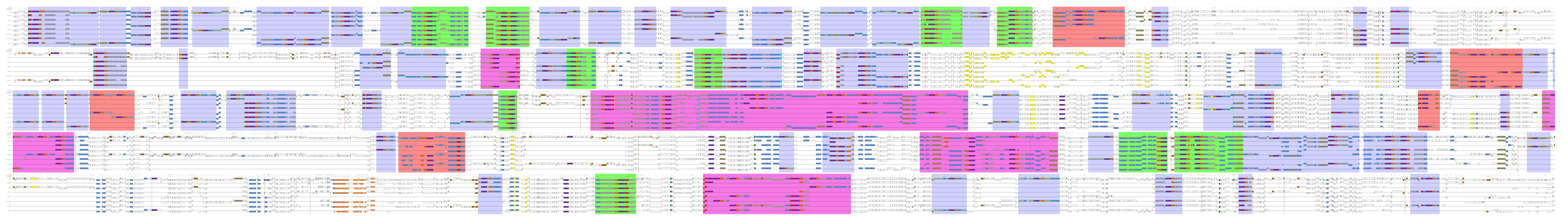

### Pudlo_Urs_et_al_Fig_S6c_pangenome4.pdf

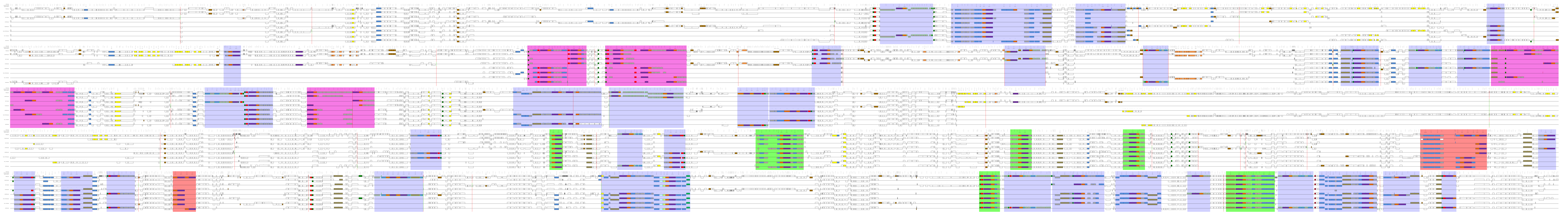
