## Supplemental Table 9 for "Phenotypic and genomic diversification in complex carbohydrate degrading human gut bacteria"

**Table S9. Supplier information for phenotypic microarray (PMA)**

| Substrate | source | 2x stock (mg/ml | sterilization |
|---|---|---|---|
| arabinose | Sigma A3131 | 10 | FS |
| cellobiose | Sigma C7252 | 10 | FS |
| fructose | Sigma F0127 | 10 | FS |
| fucose | Sigma F2252 | 10 | FS |
| galactose | Sigma G0750 | 10 | FS |
| galacturonic acid | Fluka 73960 | 10 | FS |
| glucosamine | Sigma G4875 | 10 | FS |
| glucose | Sigma G8270 | 10 | FS |
| glucuronic acid | Sigma G8645 | 10 | FS |
| mannose | Sigma M4625 | 10 | FS |
| N-acetylgalactosamine | Sigma A2795 | 10 | FS |
| N-acetylglucosamine | Sigma A3286 | 10 | FS |
| N-acetylneuraminic acid (IV-S) | Sigma A0812 | 10 | FS |
| rhamnose | Sigma R3875 | 10 | FS |
| xylose | Sigma X3877 | 10 | FS |
| chondroitin sulfate (bovine trachea) | Sigma C9819 | 10 | AC |
| heparin (porcine mucosa) | Sigma H0777 | 10 | FS |
| hyaluronan (rooster comb) | Sigma H5388 | 10 | AC |
| $\alpha$-mannan (*S. cerevisiae*) | Sigma M7504 | 10 | AC |
| neutral mucin O-glycans (porcine mucosa) | custom (porcine gastric mucosa) | 20 | FS |
| lichenan | Megazyme | 10 | AC |
| rhamnogalacturonan II | custom (red wine) | 15 | AC |
| laminarin | Sigma L9634 | 10 | AC |
| dextran (*L. mesenteroides*) | Sigma D5251 | 10 | AC |
| arabinan (sugar beet) | Megazyme P-ARAB | 10 | AC |
| arabinogalactan (larch) | Megazyme P-ARGAL | 10 | AC |
| pectic galactan (potato) | Megazyme P-GAPT | 10 | AC |
| homogalacturonan (citrus peel) | Megazyme P-GACT | 10 | AC |
| rhamnogalacturonan I (potato pectin) | Megazyme P-RHAM1 | 20 | AC |
| arabinoxylan (wheat) | Megazyme P-WAXYL | 10 | AC |
| $\beta$-glucan (barley) | Megazyme P-BGBL | 10 | AC |
| galactomannan (carob) | Megazyme P-GALML | 10 | AC |
| glucomannan (konjac) | Megazyme P-GLCML | 10 | AC |
| xylan, water soluble (oat spelt) | Fluka 95590 | 10 | AC |
| xyloglucan (tamarind) | Megazyme P-XYGLN | 10 | AC |
| inulin (chicory) | Sigma I2255 | 10 | AC |
| levan (*E. herbicola*) | Sigma L8647 | 10 | AC |
| pullulan (*A. pullulans*) | Sigma P4516 | 10 | AC |
| Glycogen | sigma G0885 | 10 | AC |
| amylopectin (maize) | Sigma 10120 | 10 | AC |
| amylopectin (potato) | Sigma 10118 | 10 | AC |

Carbohydrate sterilization methods were either by filter sterilization through a 0.22$\mu$ filter (FS);
or by autoclaving for 15 min. at 121 degrees C (AC).
